# Flow-sensitive HEG1 controls eNOS activity to prevent endothelial dysfunction, hypertension, and atherosclerosis

**DOI:** 10.1101/2025.06.24.660441

**Authors:** Michael D. Clark, Yerin Kim, Cesar A. Romero, Dong Won Kang, Kyung In Baek, Eun Ju Song, Cailin E. Kellum, Jay A. Bowman-Kirigin, Christian Park, Vir Kapoor, Jennifer S. Pollock, Hanjoong Jo

## Abstract

Hypertension (HTN), the chronic elevation of blood pressure, accounts for more atherosclerotic cardiovascular disease deaths than any other modifiable risk factor.^1^ In the arteries, stable blood flow (s-flow) drives healthy, atheroprotective endothelial cell (EC) functions including nitric oxide (NO) production, barrier function, and anti-inflammatory programs via the action of flow-sensitive proteins. We showed that s-flow stimulates Heart-of-Glass 1 (HEG1) protein expression, localization to cell-cell junctions, and secretion from ECs.^2^ We found that conditional, endothelial cell-specific knockout of (*Heg1*^ECKO^) exacerbates atherosclerosis^2^, however the mechanism was unknown. Here, we report a new role of HEG1 in controlling EC dysfunction, hypertension and atherosclerosis. We discover a novel mechanism: HEG1 regulates NO bioavailability via a flow-dependent HEG1-eNOS interaction (endothelial nitric oxide synthase, NOS3). *Heg1*^ECKO^ develops spontaneous hypertension and severe atherosclerosis, both of which are effectively treated by Angiotensin-Converting Enzyme inhibition (ACEi). UK BioBank and Swedish cohort studies reveal that plasma HEG1 levels are associated with hypertension and cardiovascular disease risk.^3,4^ Our findings suggest HEG1 may serve as a biomarker to advance personalized therapies for EC dysfunction, hypertension, and atherosclerosis.

## Introduction, Results, and Discussion

Here we tested the hypothesis that HEG1 transduces stable flow to regulate eNOS and NO bioavailability, preventing EC dysfunction, hypertension and atherosclerosis. Male and female *Heg1*^ECKO^ mice develop hypertension after tamoxifen-induced knockout, both under normal cholesterol conditions, and hypercholesterolemia induced by AAV-PCSK9 injection and Western Diet (WD) (**Figure Ai-iv**). Telemetry blood pressure measurements reveal systolic and diastolic hypertension throughout the circadian cycle.

**Figure.**
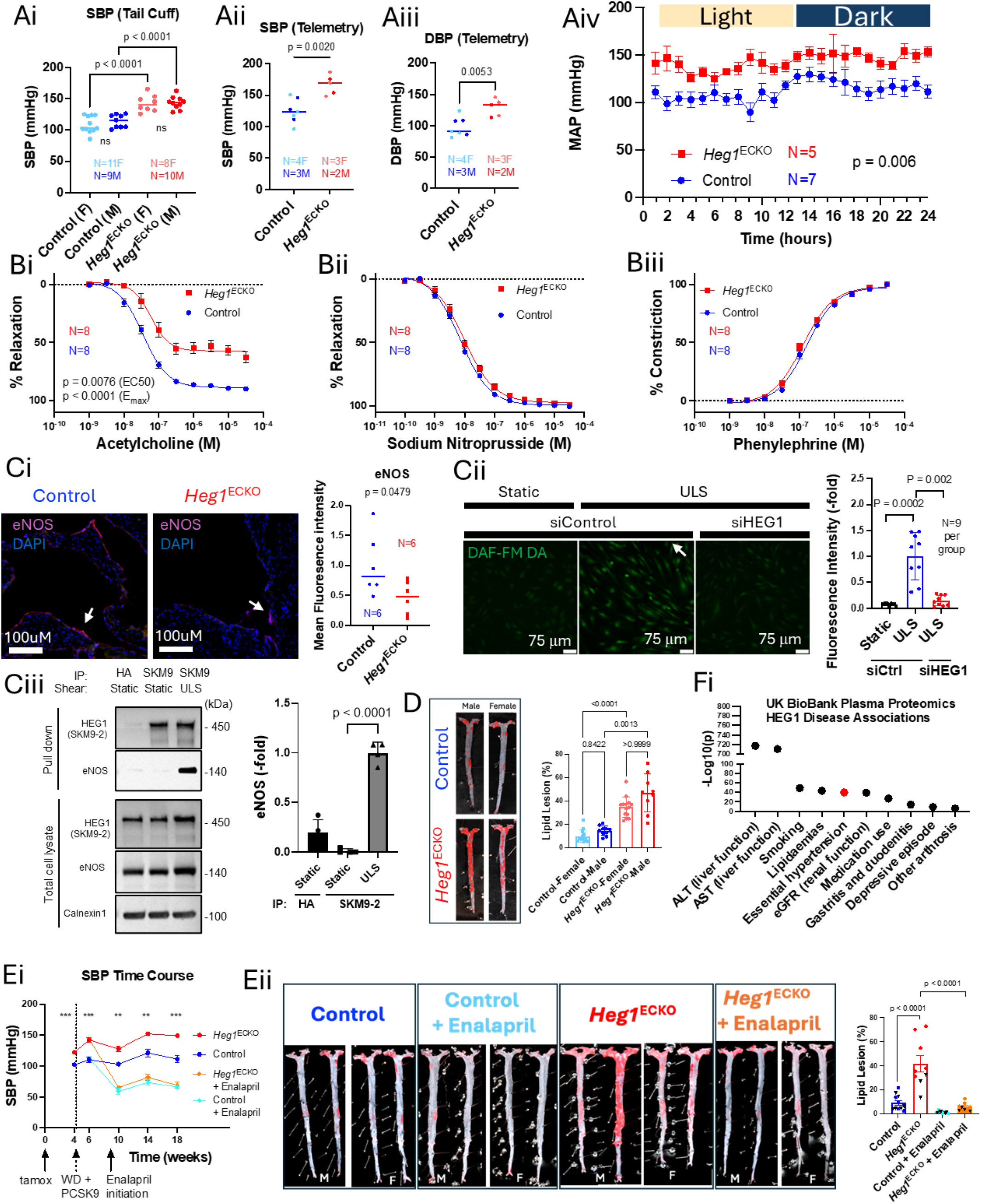
HEG1 protects against hypertension, and atherosclerosis. **Ai**. Systolic blood pressure (SBP) under hypercholesterolemia (tail cuff plethysmography, N=18-20, Control = C57BL/6J Cdh5-iCreERT2). **Aii**. SBP and **Aiii**. Diastolic blood pressure (DBP) by telemetry (Data Sciences International PA-C10 via carotid approach, normal cholesterol conditions (chow diet); female mice displayed as lighter colored symbols). **Aiv**. Mean arterial pressure (MAP) by telemetry averaged over two consecutive 24-hour cycles. No differences were detected in heart rate between groups. **B.** Aortic force myography on both male and female mice; similar results observed for both sexes. Endothelial-dependent relaxation (**Bi**), endothelial-independent relaxation (**Bii**) and contractile function (**Biii**). N= 4 male + 4 female mice per group. **Ci**. eNOS immunofluorescence imaging of mouse aortic sinus (N=6 per group). **Cii**. siRNA knockdown of HEG1 in HAECs under unidirectional laminar shear (ULS, cone and plate viscometer), imaged with Diaminofluorescein-FM diacetate ((DAF-FM-DA) to quantify NO, N=9 per group). **Ciii**. Coimmunoprecipitation of HEG1 and eNOS under static and ULS conditions in HAECs using anti-HEG1 antibody SKM9-2, with IgG beads complexed with anti-HA antibody as a negative control and Calnexin1 as a loading control. **D**. Representative Oil Red O-stained aortas after 4 months of hypercholesterolemia (N= 23-24 per group including both sexes). **E**. Enalapril study (N=9-11 per group). **Ei**. SBP time course during 4-month enalapril treatment study (tail cuff plethysmography, p< 0.01: **, p<0.001: ***, comparing *Heg1*^ECKO^ and control groups). **Eii**. Representative Oil Red O-stained aortas +/-enalapril (male (M) and female (F) aortas separated) Female mice displayed as black triangles in quantification. **Fi**. -Log10(p value) of HEG1 associations with a subset of the “20 Most Common Diseases” (For essential hypertension: -Log_10_(p) = 39.9301, Beta 0.1313).

To determine the mechanisms of *Heg1*^ECKO^ hypertension, we performed aortic vascular relaxation experiments. *Heg1*^ECKO^ impairs endothelial-dependent relaxation (acetylcholine response), while endothelial-independent relaxation (sodium nitroprusside) and contractile function (phenylephrine) are both preserved (**Figure B**). These data suggest that HEG1 may control arterial tone in an endothelial-derived, NO-dependent manner.

Next we tested how HEG1 controls NO bioavailability. *Heg1*^ECKO^ mildly reduces eNOS protein levels in the aortic sinus by immunofluorescence (**Figure Ci**). This does not fully explain such severe EC dysfunction, hypertension and atherosclerosis, leading us to hypothesize that HEG1 directly regulates eNOS activity. Indeed, HEG1 siRNA knockdown markedly reduces s-flow-induced NO production in human aortic endothelial cells (HAECs with Diaminofluorescein-FM diacetate to quantify NO, **Figure Cii**), and eNOS-Ser1177 phosphorylation with only ∼25% reduction in eNOS protein levels (data not shown). Given this surprising post-transcriptional eNOS regulation and near-complete loss of flow-induced NO production, we tested whether HEG1 and eNOS physically interact. Intriguingly, HEG1 binds eNOS specifically under s-flow conditions (**Figure Ciii**), suggesting the formation of a novel, flow-dependent HEG1-eNOS protein complex and a mechanism for direct HEG1 regulation of Enos activity. Thus, HEG1 regulates NO bioavailability at the transcriptional level (via KLF2/4)^2^ and at the novel post-transcriptional level, consistent with the severe *Heg1*^ECKO^ phenotypes.

After four months of hypercholesterolemia, male and female *Heg1*^ECKO^ mice develop severe atherosclerosis compared to controls (**Figure Di**). We next asked: to what degree does hypertension treatment reduce atherosclerosis in *Heg1*^ECKO^? ACEi enalapril decreased blood pressure and reduced atherosclerosis to below that of the control mice, completely abrogating the *Heg1*^ECKO^ effect, (**Figure Eii**) without altering blood lipid levels (not shown). This supports the central roles of hypertension and EC dysfunction in the *Heg1*^ECKO^ model of atherosclerosis.

Analysis of publicly available UK BioBank Plasma Proteomics data reveals a strong correlation between human HEG1 plasma levels and essential hypertension (**Figure Fi**).^4^ HEG1 is in the top 10% of most strongly correlated plasma protein markers of essential hypertension (not shown)^4^. Interestingly, s-flow triggers HEG1 secretion from ECs^2^, and low plasma levels of HEG1 correlate with hypertension and Framingham risk score, a clinical predictor of cardiovascular disease risk, in a Swedish cohort study.^3,4^

This study provides a novel link between HEG1, hypertension, and atherosclerosis via endothelial dysfunction, including the discovery of a novel flow-dependent HEG1-eNOS protein complex and a critical role of HEG1 in controlling NO bioavailability. Successful hypertension treatment must also prevent atherosclerosis. The robust athero-protective effects of ACE-inhibition in this HEG1-deficient model of hypertension and EC dysfunction may inform personalized therapy for patients with hypertension and EC dysfunction. While the functional role of circulating/plasma HEG1 is unknown, HEG1 may serve as a biomarker for hypertension and EC function given its plasma associations with cardiovascular disease, and identify patients most likely to benefit from ACE-inhibition.^3,4^ HEG1 may serve as a therapeutic target to mimic the antihypertensive and atheroprotective effects of s-flow, reducing residual risk factors for atherosclerosis.

Finally, we address a mystery surrounding HEG1 in rare versus common vascular diseases. HEG1 was previously linked with rare developmental disease via association with cerebral cavernous malformation (CCM) proteins.^5^ Yet, HEG1^-/-^ mice do not develop CCMs, and HEG1 variants are absent in patients with the rare CCM disease.^5^ We demonstrate the importance of HEG1 in two common cardiovascular diseases: atherosclerosis and hypertension.

## Acknowledgments

MDC, YK, CAR, JSP and HJ conceptualized and planned the experiments. All authors except JSP and HJ performed the experiments. All authors analyzed experimental results. MDC and HJ wrote and edited the final manuscript with input and approval from all authors. The authors are grateful to Dr. Ralf Adams for providing tamoxifen-inducible, endothelial specific Cdh5-iCreERT2 mice. Microscopy data for this study were acquired and/ or analyzed in the Microscopy in Medicine Core.

## Sources of Funding

This work was supported by funding from NIH grants HL119798, HL139757, and HL151358 for HJ. HJ was also supported by Wallace H. Coulter Distinguished Faculty Chair endowment. MDC and JBK were supported by NIH grant 5T32HL007745. CP was supported by NIH grant F31HL176148 and T32HL166146. KB was supported by NIH grants T32HL007745 and F32HL167625. YK was supported by AHA grant 24POST1198920.

## Disclosures

HJ is the founder of Flokines Pharma. All other authors report no conflicts of interest.

## Data Availability Statement

All data will be provided upon reasonable request.

## Methods

### Animal Studies

Animal studies presented here were approved by Emory University Institutional Animal Care and Use Committee in accordance with NIH policies. Mice were maintained in accordance with NIH guidelines in a controlled environment (21±2 °C, 50±10% relative humidity, 12-h light/dark cycle, with lights on beginning at 7:00 am Eastern Standard Time). Both male and female mice (C57BL/6J background) were included in an approximately equal balance. Control mice refer to C57BL/6J Cdh5-iCreERT2, a tamoxifen-inducible endothelial cell-specific Cre line. We generated and validated the HEG1^ECKO^ mouse line previously.^2^ Briefly, heterozygous *loxP*-flanked (floxed) HEG1 C57Bl/6J mice (HEG1^fl/+^) were generated in collaboration with Cyagen Biosciences. Homozygous floxed HEG1 mice (HEG1^fl/fl^) were subsequently developed and crossed with tamoxifen-inducible, endothelial-specific Cdh5-iCreERT2 mice. Mice were screened and genotyped by PCR and DNA sequencing. HEG1^ECKO^ was induced by intraperitoneal injection of tamoxifen dissolved in corn oil (75 mg/Kg body weight per day for 5 consecutive days).

### Chronic Atherosclerosis Studies

Hypercholesterolemic conditions were induced using AAV-PCSK9-gain of function (1×10^11^ viral genome) (Vector Biolabs #AAV8-D377Y-mPCSK9) via tail-vein injection for low-density lipoprotein receptor (LDLR) knockout, combined with *ad lib* high-fat Western Diet feeding (Teklad, Inotiv TD.88137).^2^ Plaque quantification was performed as described previously.^2^ Briefly, the mice were sacrificed and the entire aortic tree excised and gross macroscopic images were collected after *en face* Oil Red O staining (Sigma-Aldrich No. MAK194). Serum cholesterol and triglyceride concentrations were determined using a Beckman CX7 biochemical analyzer at the Cardiovascular Specialty Laboratories, as reported previously.^2^ The chronic atherosclerosis study shown in Figure panel **D** comprised 5 months of inducible HEG1 knockout, with the last 4 months under hypercholesterolemic conditions.

### Enalapril Study

Enalapril Maleate (Med Chem Express. HY-B0331A) was administered to mice in the drinking water at a final dose of 40 mg/L drinking water, following a run-in period of 120 mg/L and 80 mg/L during which the optimal dose was determined. Drinking water with enalapril was changed twice per week. For the enalapril study, the time course is shown in Figure panel **Ei**, and mice were sacrificed following the final 18 week blood pressure measurement. Aortas were harvested and lipid lesions were quantified with Oil Red O as described above and previously.^2^

### Non-Invasive Blood Pressure Measurements

Tail cuff plethysmography was performed using a BP-2000 series II Blood Pressure Analysis System (Visitech Systems) as we have done previously.^6^ Measurements were taken at a consistent time of day for all mice (afternoon), and mice were trained for multiple sessions prior to the initial blood pressure readings. For each day of blood pressure measurements, 10 measurements were obtained and averaged for each mouse. Data shown in Figure Panel **Ai** are Systolic blood pressures (SBP) collected after two months of inducible HEG1 knockout, the final one month of which were under hypercholesterolemic conditions.

### Telemetry Blood Pressure Measurements

Telemetry was performed as we have done previously, with minor modifications described as follows.^7^ Mice at ∼3.5 months of age (subjected to ∼10 weeks of inducible HEG1 knockout, on chow diet) were implanted with PA-C10 telemeters (Data Science International) in the carotid artery to obtain intra-arterial blood pressure measurements in ambulatory mice. After 1 week post surgical recovery period, blood pressure and heart rate parameters were recorded every 20 minutes for 2 minutes. Data were collected over two consecutive 24 hour cycles and averaged to generate data shown in Figure panels **Aii**-**Aiv**. Telemetry studies were conducted under normal cholesterol conditions.

### Force Myography Measurements

Force myography experiments were conducted as we did previously, with minor modifications described as follows.^6^ Briefly, mice at 3-3.5 months of age under normal cholesterol conditions were sacrificed and their thoracic aortas collected without perfusion. Aortas were cleaned of surrounding fat, and were trimmed to ∼2mm length rings, which were then mounted on pins in a DMT 620M force myograph (Danish Myo Technology). Aortas were maintained in physiological saline solution (PSS) and chambers continuously bubbled with a mixture 95% O2 5% CO2 at 37°C. Aortas were equilibrated and brought to a resting tension of 10 millinewton (mN). Aortas were then subjected to integrity testing using phenylephrine pre-constriction with a single-dose acetylcholine response. For relaxation curve experiments, aortas were pre-constricted with 1 micromolar phenylephrine (Med Chem Express HY-B0471) and relaxed with increasing concentrations of acetylcholine (Sigma Aldrich A6625) or sodium nitroprusside (Sigma Aldrich 71778). PSS buffer contains 130 mM NaCl, 4.7 mM KCl, 1.18 mM KH_2_PO_4_, 1.17 mM MgCl_2_, 14.9 mM NaHCO_3_, 5.5 mM Glucose, 0.026 mM EDTA, and 1.6 mM CalCl_2_.

### Immunofluorescence Imaging of Mouse Aortic Sinus

Control and HEG1^ECKO^ mice were subjected to 5 months of inducible HEG1 knockout, the last 4 months of which were under hypercholesterolemic conditions. Immunofluorescence imaging was performed as previous described.^2^ Briefly, mice were sacrificed and the hearts including the aortic sinus and root and fixed in 10% formalin for 24 hours. The aortic sinus was isolated and embedded in Optimal Cutting Temperature (OCT, Sakura #4583) media, frozen and cryosectioned. Samples were cryosectioned and stained for eNOS (primary antibody: eNOS (D6A5L) Rabbit mAb Cell Signaling #32027S at 1:100, and secondary antibody Donkey anti-rabbit IgG Alexa 647, Invitrogen #A-31573). Samples were DAPI-mounted for fluorescence imaging on an inverted fluorescence microscope (Keyence #BZ-X800, Japan).

### Endothelial Cell Culture, Shear Exposure, Short Interfering RNA Knockdown, and Nitric Oxide Bioavailability Measurements

Primary human aortic endothelial cells (HAECs; Cell Applications) were cultured as we have described previously.^2^ HAECs were cultured in a complete medium consisting of MCDB 131 (Corning No. 15-100-CV), 10% fetal bovine serum (R&D Systems No. S11550), 1% penicillin-streptomycin (Gibco No. 15140-122), 1% L-glutamine (Gibco No. 25030-081), 1% EC growth supplement from bovine brain extract, 50 μg/mL of L-ascorbic acid (Sigma No. A5960), 1 μg/mL of hydrocortisone (Sigma-Aldrich No. H088), 10 ng/mL of epidermal growth factor (EGF; STEMCELL Technologies No. 78006), 2 ng/mL of insulin-like growth factor 1 (IGF-1; R&D Systems No. 291-G1), 2 ng/mL of fibroblast growth factor (FGF; ProSpec No. cyt-218-b), and 1 ng/mL of vascular endothelial growth factor (VEGF; BioLegend No. 583706). HAECs were cultured on gelatin-coated (Sigma No. G1890) dishes and used for experimentation between passages 4 and 8.

Shear exposure was performed using the cone and plate viscometer as we have done previously.^2^ HAECs grown to confluency on gelatin-coated 100-mm tissue culture dishes (Falcon No. 353003) were exposed to ULS (+15 dyn/cm^2^) mimicking s-flow for 24 hours.

Short interfering RNA (siRNA) knockdown was performed as we previously described, using oligofectamine (Invitrogen No. 12252-011) transfection agent and the same siRNA sequences (for HEG1 and control) listed in our prior publication.^2^

Nitric Oxide measurements in siRNA-treated HAECs in response to shear stress were performed using 4-amino-5-methylamino-2’,7’-difluororescein diacetate (DAF-FM DA, Med Chem Express HY-D0717). HAECs in basal media (MCDB131 + 0.5% fetal bovine serum) were subjected to 24 hours of ULS and then incubated with 1uM DAF-FM DA at 37°C for 30 min. Following two phosphate buffered saline washes, HAECs were then fixed in 2% paraformaldehyde (5 min at 4°C), washed and imaged on a confocal fluorescence microscope (Leica Thunder, Excitation max 488nm, and Emission max 520nm). Fluorescence intensity was quantified in ImageJ to determine NO bioavailability.

### HEG1 eNOS Coimmunoprecipitation Experiments

HAECs were subjected to 24 hours of ULS were washed, harvested and lysed using Pierce IP Lysis buffer (Thermo Fisher Scientific No. 87787) as we previously described.^2^ Lysates (1.2 mg/ml total protein concentation) were pre-cleared with IgG beads for 1 hour, followed by overnight incubation with either 5 µg of anti-HEG1 SKM9-2 mouse monoclonal IgG (provided by Dr. Shoutaro Tsuji)^2^ or anti-HA mouse monoclonal IgG (negative control, Invitrogen 26183) per lysate. Samples were then incubated for 3 hours with 20ul of protein A/G magnetic beads (Pierce VWR 88803).

SDS-PAGE and Western Blot were peformed as we described previously.^2^ Briefly, membranes were incubated at 4 °C overnight with the following primary antibodies: anti-calnexin rabbit polyclonal antibody (1:2000; Abcam No. ab75801), anti-eNOS mouse monoclonal antibody (250 ng/mL; BD Biosciences No. 610297), and anti-HEG1 SKM9-2 mouse monoclonal antibody (5 μg/mL). Membranes were incubated at room temperature for 1 hour with the following secondary antibodies: goat anti-mouse IgG horseradish peroxidase (Cayman Chemical No. 10004302) and goat anti-rabbit IgG horseradish peroxidase (Cayman Chemical No. 10004301). Blots were incubated with Immobilon Western chemiluminescent horseradish peroxidase substrate (Millipore No. WBKLS0500) and imaged and quantified using the iBright FL1000 imaging system.

### UK BioBank Plasma Proteomics Analysis

From the publicly-available, published UK BioBank data, -Log10(P) values were plotted for the association of HEG1 plasma levels with a subset of the “Proteomic associations with smoking, medication use, liver function, renal function and top 20 most prevalent diseases”.^4^ The 10 most significant associations (as measured by P value) are shown in Figure panel **Fi**. The source data are available in the supplemental materials of the UK BioBank Plasma Proteomics publication: “Plasma proteomic associations with genetics and health in the UK Biobank”.^4^

### Statistical Analyses

Data are presented as mean + SEM. Statistical analyses were performed in GraphPad Prism v10. The statistical testing used for data in each figure panel are: Unpaired t-test (Ai-iii, Ci, Cii), Unpaired t-test with Bonferroni correction (Ei), Kruskal-Wallis test with Dunn’s multiple comparisons (Cii, D), One way ANOVA with Bonferroni (Eii), Two way ANOVA (Aiv), 4-parameter logistic model with Extra Sum of Squares F-test on fit parameters (Bi). For panels Cii and D, non-parametric methods (Kruskal-Wallis test with Dunn’s multiple comparisons) were employed based on Shapiro-Wilk normality tests, which suggested that at least one group within Cii and D were inadequately described by a normal distribution (P<0.05). Normality testing was not performed for Aiv and Ei (due to time-series data), Ci-iii (due to statistical testing on fit parameters), and Ciii (due to N=4 per group). All other data met Shapiro-Wilk criteria for normality (P>0.05).

